# Effect of Water Fluoridation on VM-M3 Tumor Metastasis

**DOI:** 10.1101/2023.06.30.547222

**Authors:** Gerald M. Mastellone, Nathan L. Ta, Purna Mukherjee, Derek C. Lee, Jack H. Maurer, Thomas N. Seyfried

## Abstract

Water fluoridation has been used for combatting dental caries in the general population. Although fluoride has been evaluated as a possible carcinogen, no studies have evaluated the influence of water fluoridation on tumor metastasis. *Ex vivo* bioluminescent imaging and tumor volume measurements were used to assess the influence of water fluoridation on the metastatic spread of VM-M3/Fluc tumor cells grown in their syngeneic inbred VM/Dk mice that received either sodium fluoride (NaF) or hydrofluorosilicic acid (HFSA) in the drinking water (at either 5.0 mg/L or 20.0 mg/L) for 35 days. During the course of the study, there was no difference in water intake or body weight between the control mice and mice that drank either NaF or HFSA. No significant differences were found for VM-M3/Fluc organ metastasis in the mice that drank water with either 5.0 mg/L or 20.0 mg/L NaF or HFSA versus control mice that drank water without fluoride. Both overall survival and flank tumor volumes were similar in control mice and in mice that drank water containing HFSA. In conclusion, the results showed that fluoride present in the drinking water had no statistically significant effect on the metastatic spread of VM-M3/Fluc tumor cells nor on mouse survival.

## Introduction

The effect of fluoride on dental health, specifically in preventing dental caries, has been studied for over a century and has revealed that exposure to low levels of fluoride reduces the incidence of dental caries (1,2). Previous research suggested that fluoridation levels of drinking water can be beneficial for human dental health, but excessive exposure can result in dental or skeletal fluorosis (3,4). However, there are concerns that excessive fluoridation of water can be carcinogenic, increasing incidence of cancer, particularly osteosarcoma in boys (5). Yet, population studies have shown no link between fluoride consumption and cancer incidence (6– 8). This topic remains controversial with strong opposing views. In this study, we sought to determine if water fluoridation could influence the metastatic spread of the VM-M3/Fluc tumor cells in the syngeneic VM/Dk inbred mouse host.

Metastasis is the term used to describe the spread of cancer cells from a primary tumor site to surrounding tissues and organs (9). When implanted into the flank of the mouse, the VM-M3/Fluc tumor cells spread to multiple organ systems including the brain, kidney, liver, spleen, and lungs (9–11). The VM-M3/Fluc tumor cell line had been stably transfected with the firefly *luciferase* gene which enables the visualization of metastasis and measurement of the degree of tumor invasion within the mouse *in vivo* or *ex vivo* via bioluminescent imaging (10). Using these techniques, tumor metastasis was compared between groups of mice receiving different quantities of fluoridation in their drinking water.

Both sodium fluoride (NaF) and hydrofluorosilicic acid (HFSA) have been used for artificial water fluoridation globally as well as fluoridation of other products such as toothpastes (12,13). Despite these similarities, HFSA is commonly preferred for water fluoridation because it is a more cost-effective chemical and a by-product of phosphate chemical production (12). In our experiments, both NaF and HFSA were used in separate, repeated trials to determine if there were differences in their effect on VM-M3/Fluc metastasis. The recommended level of fluoridation in drinking water in the United States is 0.7 – 1.2 mg/L fluoride (13). Our study compared a control of 0 mg/L fluoride with concentrations of 5.0 mg/L and 20.0 mg/L of either NaF or HFSA. These concentrations are higher than the recommended concentration of fluoridation in water for humans because mice have a basal metabolic rate that is approximately 7 times greater than that of humans (14). The supraphysiological concentration of 20 mg/L was used to exacerbate any potential effect of metastasis. Our findings in this preliminary study showed that water fluoridation had no statistically significant effect on VM-M3/Fluc tumor cell metastasis.

## Materials and Methods

### Mice

The mice used during this experiment were received as gifts from G. Carlson (McLaughlin Research Institute, Great Falls, Montana) and from H. Fraser (University of Edinburgh, Scotland). This experiment utilized the VM/Dk (VM) strain of mice housed within the Boston College Animal Care Facility as previously described (15). Both female and male mice (∼90 days of age) housed in groups of five or less by sex in plastic cages with filter tops and Sani-Chip bedding (P.J. Murphy Forest Products Corp., Montville, NJ). The mice were fed a standard chow diet *ad libitum* and drank from standard cage water bottles containing either reverse osmosis de-ionized water (RODI) alone or with 5.0 mg/L or 20.0 mg/L of NaF or HFSA. All animal procedures were conducted in accordance with the NIH Guide for the Care and Use of Laboratory Animals and were approved by the Animal Care Facility at Boston College.

### Drug preparation

Both sodium fluoride (NaF, Fisher Scientific) and hydrofluorosilicic acid (HFSA, Sigma-Aldrich) were used during separate trials of the experiment. NaF was measured by weight and mixed into the drinking water of the mice in quantities of 0 mg/L, 5.0 mg/L, and 20.0 mg/L. HFSA was purchased in a 20-25% by weight mixture. Based on our calculations, a quantity of 22.5 ul and 90.0 ul of HFSA per liter of water was equivalent to a concentration of 5.0 mg and 20.0 mg per liter of water. The HFSA in those amounts was added and mixed into the drinking water of the mice. Both compounds were mixed in glass bottles with 1.0 L of RODI water. This mixture was transferred from its container directly into the water bottles in the mouse cages used during the experiment. Fresh mixtures were prepared weekly.

### Origin of the VM-M3/Fluc tumor

The VM-M3/Fluc tumor arose spontaneously in the cerebrum of a male adult VM mouse as previously described (10). *In vivo* viability of the VM-M3/Fluc tumor was preserved through serial transplantations and the preparation of the VM-M3/Fluc cell line was previously described (10,11).

### Subcutaneous tumor implantation

The VM-M3/Fluc cells were implanted subcutaneously into a VM mouse as described previously (9). The tumor from donor mice was excised and chopped into tiny fragments. Each mouse received 0.2 mL of tumor fragments suspended in 0.3 mL of PBS. Mice were anesthetized with isofluorane (Covetrus, Ohio) and the tumor was implanted by subcutaneously using 1 cc tuberculin syringe and 18-gauge needle into the right flank.

### Tumor growth and metastasis analysis using bioluminescent imaging

The experiment consisted of a pretrial and trial period as shown in Figure 1. During the pretrial period, mice were separated into 1 of 3 groups; control (0 mg/L), 5.0 mg/L, or 20.0 mg/L NaF or HFSA in drinking water. Mice were given water *ad libitum*. The body weight of each mouse and water intake per cage was monitored closely throughout the experiment to monitor the health of the mice. After approximately fourteen days post subcutaneous tumor implantation, each mouse underwent bioluminescent *in-vivo* imaging to measure tumor growth as described (11).

**Figure 1.**
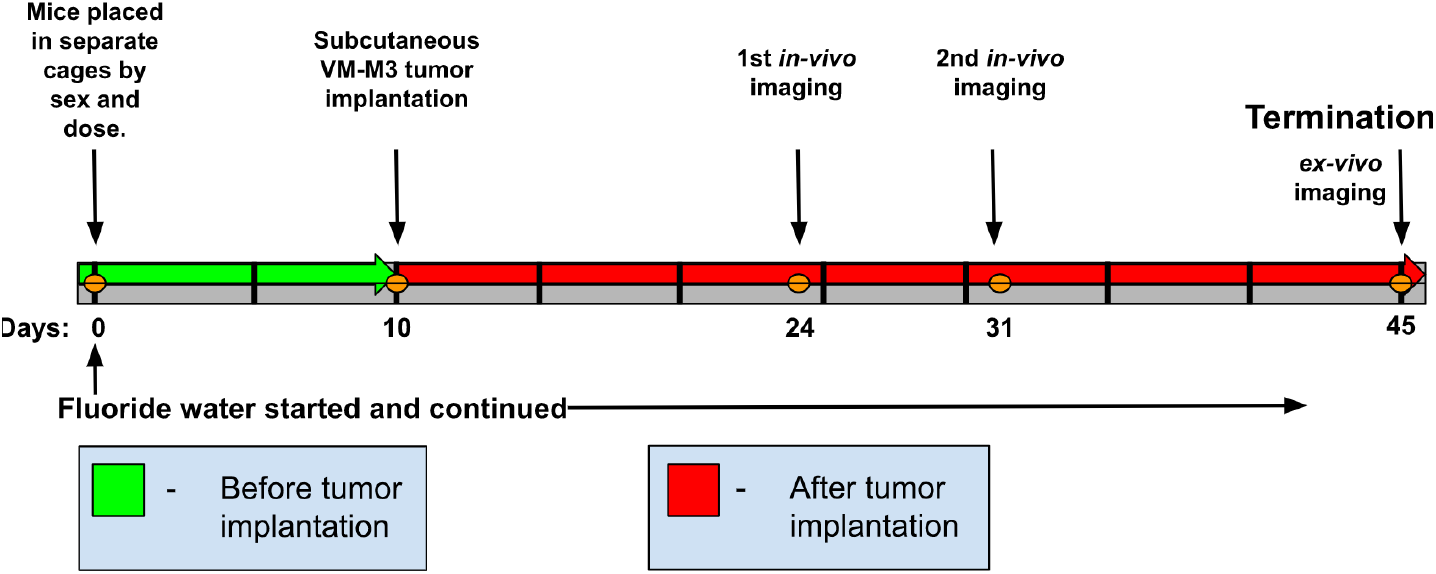
Timeline of Experiment with Days of Treatment and Imaging. The figure depicts the experimental timeline used for both the NaF and HFSA experiments. The fluoride dosages used (dose) are described in the text.

**Figure 2.**
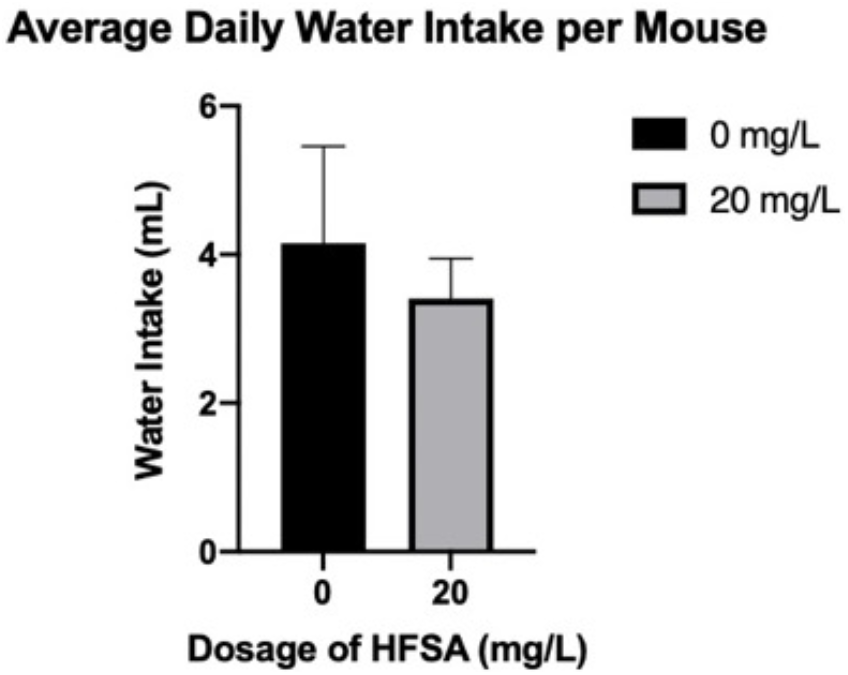
Influence of HFSA on Water Intake. The estimated daily water intake (mL) consumed per mouse in the control and HFSA groups. The water bottles were weighed 3-4 times per week to estimate daily water consumption/mouse. The change in weight was measured and averaged for the number of days that had passed and number of mice in the cage. The differences in water consumption were not statistically significant between the groups.

Mice were imaged with the Spectral Instruments Imaging AMI HTX system to measure the bioluminescent signal from the tumors. For *in vivo* imaging, mice received an i.p. injection of *D*-luciferin. Imaging occurred in durations of 5 minutes. Normal procedure and monitoring continued. After either thirty days or when the control mice reached morbidity (excess tumor burden and ulceration), the mice were then sacrificed to determine the extent of metastatic spread. For metastasis analysis, the individual organs (brain, liver, kidney, lungs, and spleen) were excised and imaged *ex vivo* for 2 mins.

### Tumor volume and survival analysis

Upon sacrifice of the VM-M3 mice at morbidity, tumors were measured. A caliper was used to measure tumor volume as calculated using (Width2 x Length) / 2.

### Survival Analysis

The VM-M3/Fluc tumor was implanted subcutaneously and used as day 0. Mice in either the control group receiving 0 mg/L of fluoride or the experimental group receiving fluoridated water of 20.0 mg/L received water *ad libitum* for approximately five weeks. During the third HFSA trial where mice did not undergo bioluminescent imaging, individual mice were sacrificed at morbidity.

### Statistical Analysis

Bioluminescence levels were recorded and compared by organ between the experimental and control groups using the nonparametric Wilcoxon test due to the small sample size of each experiment. For bioluminescent values to be considered significant, they were compared to bioluminescent values of non-tumor bearing VM-M3 mouse bioluminescent values for each organ. The results of the statistical tests were then placed on a box and whisker plot to compare means and standard deviations visually as shown in Figures 3-4. Survival study was plotted on a Kaplan-Meier curve using Graphpad Prism software and significance was determined by a log-rank test.

**Figure 3.**
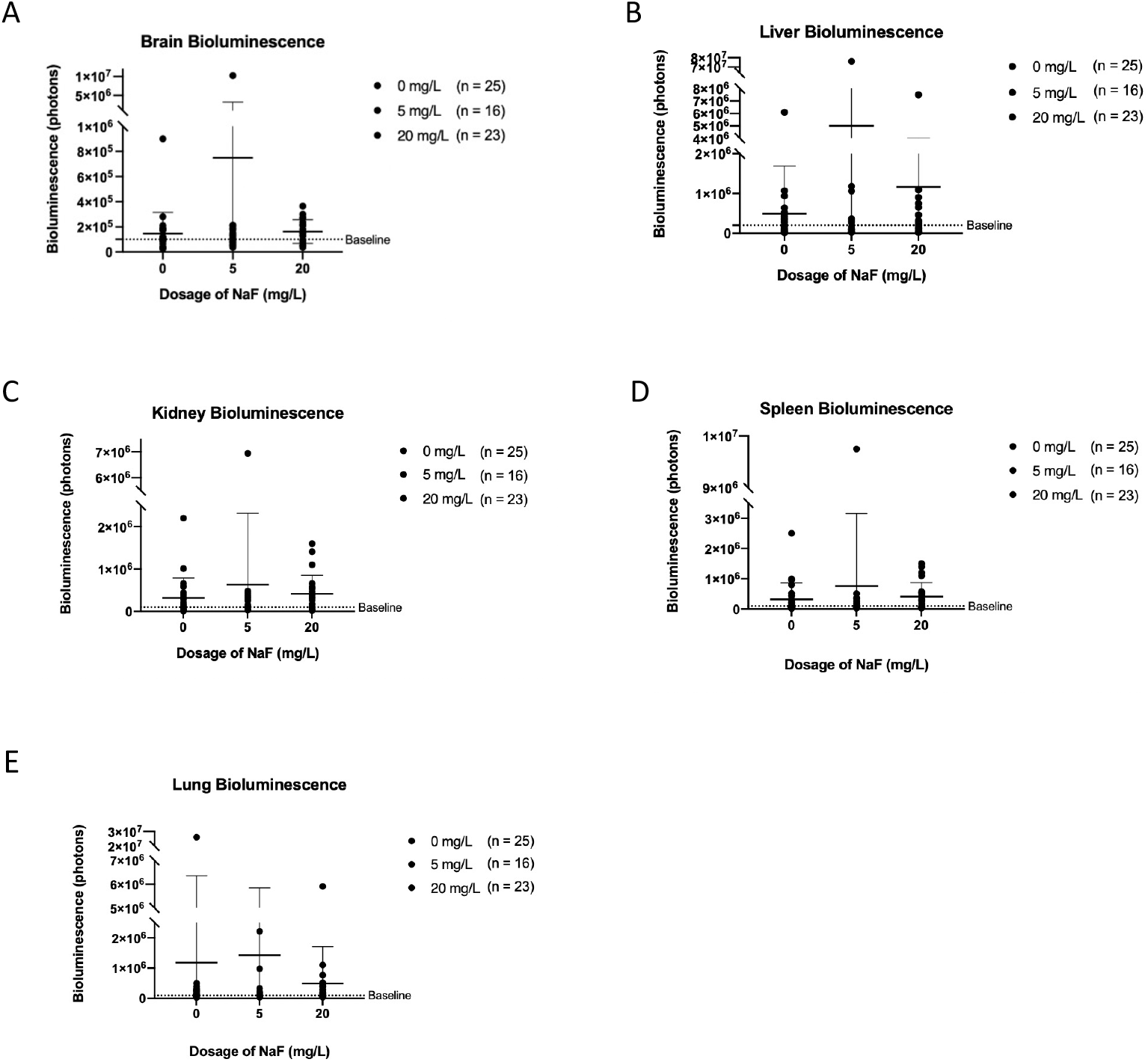
Influence of NaF on Organ Metastasis. The bioluminescence of brain (A), liver (B), kidney (C), spleen (D), and lung (E) between the control and treated groups, in either 5.0 mg/L or 20.0 mg/L are depicted. The data represent the combined values from 4 separate trials. The y-axis shows the quantified amount of bioluminescence in each respective organ where each dot represents one mouse. The dotted line termed the “baseline” at 1.0 × 105 photons of bioluminescence represents the organ bioluminescence value in non-tumor bearing mice. No significant differences were found for organ bioluminescence between mice that drank unfluoridated water and those that drank fluoridated water at either NaF concentration.

## Results

The objective of this study was to determine if water fluoridation could influence the metastatic spread of VM-M3/Fluc cells in their inbred syngeneic VM/Dk mouse host strain. The study was not designed to test the influence of water fluoridation on the incidence of cancer in these mice. Two doses of NaF or HFSA (5.0 mg/L or 20.0 mg/L) were used in the experiments. As mentioned previously, mice have a metabolic rate that is approximately 7x greater than that of humans (14), hence 5.0 mg/L concentration was used (recommended concentration of 0.7 mg/L x 7). Due to a relatively short treatment exposure time (about 45 days including pre-treatment), the higher supra-physiological concentration of 20 mg/L was used to increase the concentration of fluoride in the mice. Bucher *et al*. showed that increased bone fluoride content in mice and rats was correlated with the concentration of their fluoride intake (16). This could provide some insight as to whether prolonged consumption of fluoridated water in humans might influence metastasis in already formed tumors.

### Water Intake and Body Weight

The average water intake per mouse between the control group and 20.0 mg/L group of the HFSA trial showed no statistically significant differences (Figure 2). Similarly, during the NaF trials (data not shown), water intake was measured for several days throughout each trial and no noticeable difference in average water intake per mouse was noticed between groups.

### Influence of Water Fluoridation on Organ Metastasis

The bioluminescent imaging technique used to determine levels of tumor metastasis was described previously in the Materials and Methods section. The data shown in Figure 3 and Figure 4 represent the bioluminescence values in both NaF and HFSA trials, respectively. Results from Figure 3 were data combined from 4 individual trials. Fluoridation of water using NaF resulted in no significant differences in bioluminescence in the brain, liver, kidney, spleen, and lung between the control, 5.0 mg/L, or 20.0 mg/L groups (Figure 3 A-E).

**Figure 4.**
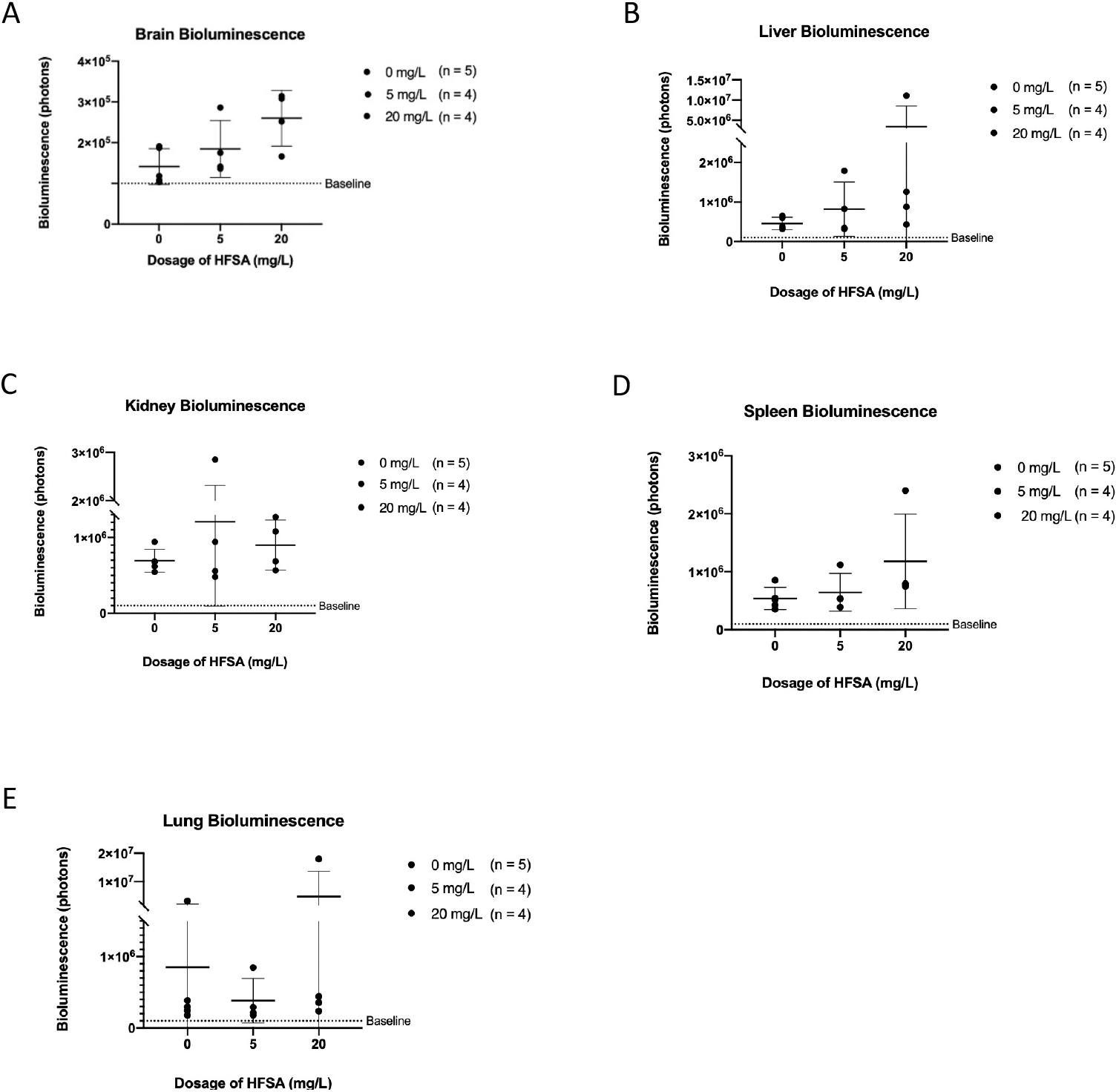
Influence of HFSA on Organ Metastasis. The bioluminescence of brain (A), liver (B), kidney (C), spleen (D), and lung (E) between the control and treated groups, in either the 5 mg/L or the 20 mg/L are depicted. The y-axis shows the quantified amount of bioluminescence in each respective organ where each dot represents one mouse. The dotted line termed the “baseline” at 1.0 × 105 photons of bioluminescence represents the organ bioluminescence value in non-tumor bearing mice. No significant differences were found for organ bioluminescence between mice that drank unfluoridated water and those that drank HFSA fluoridated water.

Results from Figure 4 were from the 2nd experimental trial. A preliminary study (data not included), carried out using only the higher and supra-physiological concentration of 20 mg/L HFSA, resulted in a trend of higher metastasis, but was not statistically significant. Hence, in the second experimental trial, data shown in Figure 4, both 5.0 mg/L and 20.0 mg/L concentrations were tested. Fluoridation of water using HFSA resulted in no significant differences in bioluminescence in the brain, liver, kidney, spleen, and lung between the control and treated groups, in either 5.0 mg/L or 20.0 mg/L of HFSA (Figure 4 A-E). Similar to our preliminary study, we did observe a trend of higher metastasis in the 20.0 mg/L of HFSA mice, however the results were not statistically significant.

### Tumor Volume

Tumor volume was recorded for the 2nd experimental trial using HFSA and measured as previously described in the Materials and Methods. Fluoridation of water using HFSA yielded no significant difference in tumor volume between the control and both the treated groups (Figure 5).

**Figure 5.**
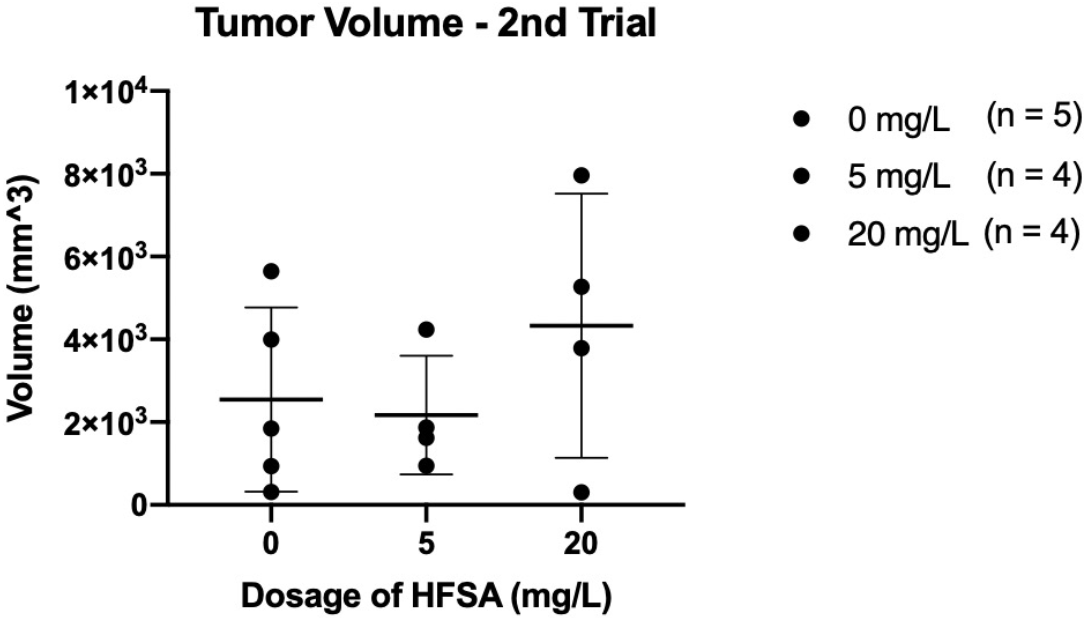
Influence of HFSA on VM-M3 Tumor Volume. The graph show tumor volumes as measured at the time of death for each mouse. The y-axis describes the tumor volume in mm3 where each dot represents each individual mouse. There was no signicant difference was found for tumor volumes between the control and both the treated groups.

### Influence of HFSA Water Fluoridation on Mouse Survival

The log-rank statistical analysis test showed no significant difference between the control and the HFSA treated mouse group (Figure 6). In this experiment, the mice received HFSA only at the 20.0 mg/L concentration since an effect on survival would likely be seen at a higher than at a lower HFSA concentration. All mice died between 19-22 days post-tumor implantation.

**Figure 6.**
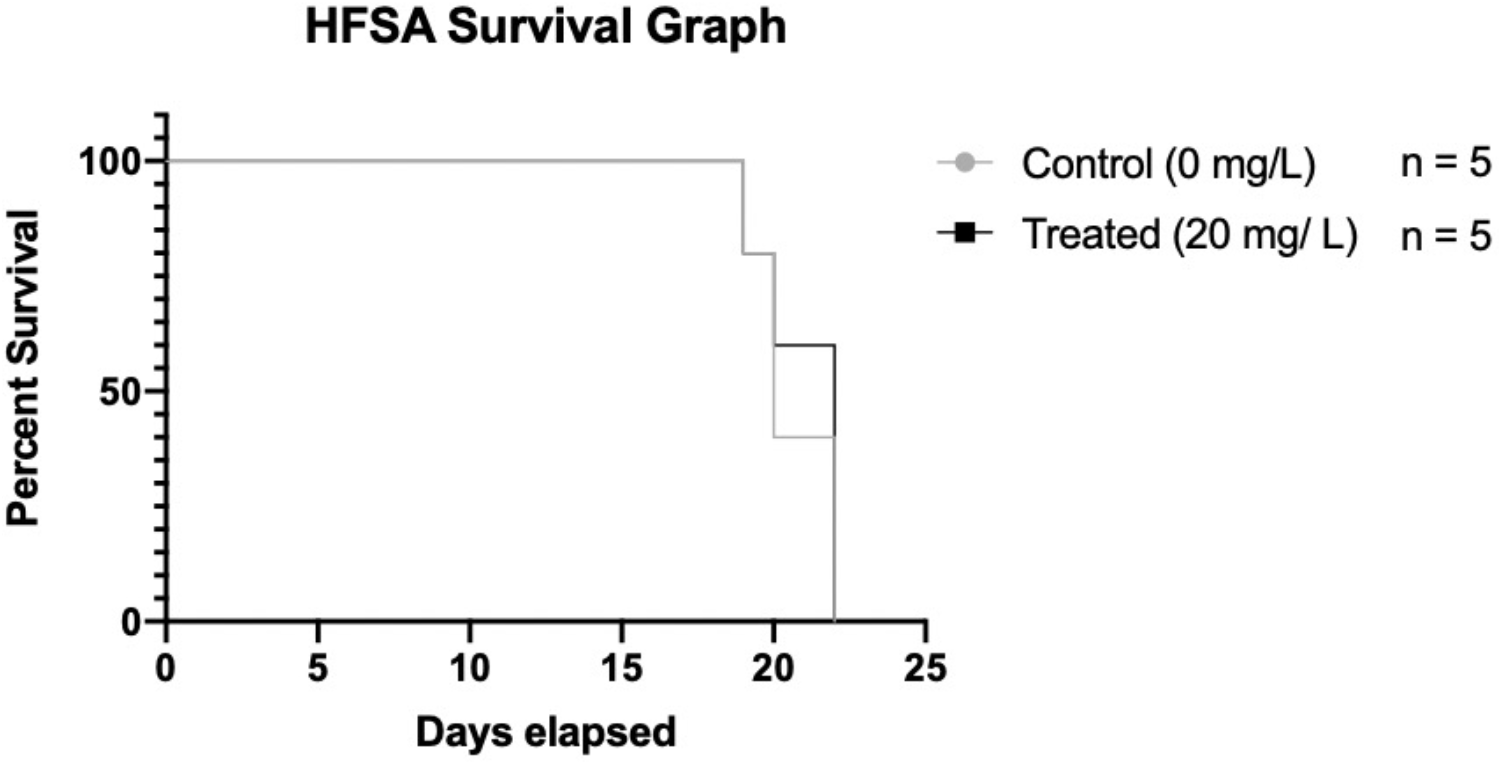
Influence of HFSA on Survival. A Kaplan–Meier survival plot was used to measure mouse survival. The log-rank statistical analysis showed no significant difference between the control and the treated group. No mouse in either group survived past 22 days post-implantation during this trial.

## Discussion

The topic of fluoridation of municipal drinking water is controversial. On one hand, studies including the US National Toxicology Program concluded that fluoride did not increase the incidence of carcinogenesis in either male or female rats and mice over a two-year period (16). The results of this study showed that 4 of 130 male rats developed bone cancer after drinking a high concentration of fluoride at either 100 or 175 ppm (1ppm = 0.998 mg/L). These fluoride concentrations were nearly 5 to 35 times higher than the fluoride concentrations used in our study and were well beyond the physiological level present in fluoridated drinking water in the U.S. and abroad (16). On the other hand, Bassin *et al*., reported a higher incidence of osteosarcoma in young male children from higher exposure to fluoride (5). Population studies at municipalities and national levels, including US, Ireland, and Taiwan were unable to find significant correlations in cancer incidence and mortality in humans that consumed fluoridated water (6–8). Hence, the linkage of water fluoridation and carcinogenesis remains ambiguous.

No prior studies have evaluated a linkage of water fluoridation to the metastasis of already developed cancers. We evaluated this linkage using the highly metastatic VM-M3 tumor cells grown in their syngeneic inbred VM/Dk mouse host (9-11). No statistically significant differences were found between the metastatic spread of VM-M3 tumor cells between VM/Dk mice that did or did not drink fluoridated water. Although there appeared to be an upward trend in metastatic spread and tumor volume at the higher unphysiological dose of HFSA (20 mg/L), the difference was not statistically significant nor was overall mouse survival. Our studies do not rule out the possibility that HFSA might influence metastasis in the VM-M3 model if used at even higher concentrations or if evaluated in other models of metastasis, but more extensive studies with larger sample sizes would be needed to support this possibility.

## Abbreviations

DMEM: Dulbecco Modified Eagle Medium
FBS: fetal bovine serum
HFSA: hydrofluorosilicic acid
i.p.: intraperitoneal
NaF: sodium fluoride
PBS: phosphate buffered saline
RODI: reverse osmosis de-ionized water

